# Highly accurate prophage island detection with PIDE

**DOI:** 10.1101/2025.02.26.640259

**Authors:** Hongyan Gao, Bowen Li, Zihan Guo, Lei Zheng, Junnan Chen, Guanxiang Liang

## Abstract

As important mobile elements in prokaryotes, prophages shape the genomic context of their hosts and regulate the structure of bacterial populations. However, it is challenging to precisely identify prophages through computational methods. Here, we introduce PIDE for identifying prophages from bacterial genomes or metagenome-assembled genomes. PIDE integrates a pre-trained protein language model and gene density clustering algorithm to distinguish prophages. Benchmarking on bacterial genomes with experimental prophage annotation demonstrates that PIDE pinpoints prophages with precise boundaries. Applying PIDE to 4,744 human gut representative genomes reveals 24,467 prophages with widespread functional capacity. PIDE is open source and is available at https://github.com/chyghy/PIDE.

## Background

Bacteriophages, or phages, are viruses that infect bacteria and the most abundant and diverse entities in the biosphere(1). They mainly exhibit two life cycles: the lytic cycle and the lysogenic cycle. Unlike obligate lytic phages, temperate phages have a life cycle that involves a decision-making process between lysis and lysogeny. During lysogeny, temperate phages integrate their genomes into bacterial genomes as prophages, remaining dormant until an induction signal is sensed(2). A previous study found that over 80% of bacterial strains from infant feces could produce phages when exposed to mitomycin C(3), highlighting the prevalence of lysogeny in the human gut. As a result, lysogeny not only regulates bacterial community structure but also contributes to genetic variation by disrupting host genes or introducing new functions(4–7), and certain defective prophages can have a significant impact on the host’s physiological functions(8–10). Therefore, identifying prophages, even defective prophages, in a given environment is essential for understanding their effects on bacterial hosts. Due to the difficulty in determining the activity status of prophages, this study uses the term "Prophage Island (PI)" to refer to phage-related genomic islands.

Given their significance, different prophage identification tools have been developed, such as Virsorter2, PHASTER, Prophinder, ProphET, PhiSpy and the recently published geNomad(11–17). One of the most common strategies for predicting prophages is identifying phage-like Open Reading Frames (ORFs) based on sequence similarity searches or machine learning classifiers, and subsequently detecting regions with a high enrichment of phage ORFs. The success of pre-trained language models in natural language processing has spurred a growing interest in applying similar deep learning techniques to biological tasks. Protein language models based on Transformer architecture are adept at uncovering complex patterns within protein sequences to decipher their structures and functions. Evolutionary Scale Modeling (ESM)-2, a protein language model pre-trained on millions of unique protein sequences from the UniRef database, can capture both sequence information and functional characteristics of proteins(18). It can be easily adapted by finetuning a small proportion of parameters for specific tasks(19,20).

Here we introduce a tool named PIDE (Prophage Island Detection using ESM-2) to detect prophage islands. PIDE first accurately predicted phage ORFs from bacterial genomes by finetuned ESM-2 with 650 million parameters, and subsequently pinpointed prophages by employing a statistical algorithm to cluster predicted phage ORFs based on gene density. Further, we verified the performance of PIDE using the experimental data. Last, we applied it to bacterial genomes derived from human gut microbiome data and found that PIs were widely present in gut bacteria and functional analysis indicated that these PIs could potentially influence their host resistance and metabolism.

## Results

### The overview of the PIDE framework

PIDE is an automated, user-friendly tool specifically developed for the identification of PIs within bacterial genomes. The tool consists of two core stages: (1) identifying phage ORFs; (2) detecting regions with high enrichment of these ORFs. In the first step, the deep learning model for phage ORF prediction adopts ESM-2 with 650 million parameters as the extractor for 1D sequence embedding (Fig.1a). Then the sequence embedding is fed into a multi-layer perceptron (MLP) classifier with five layers to obtain the classification for each ORF, where ORFs with the probability over 0.5 are considered as phage ORFs. Next, as many PIs may carry bacterial genes which are also known as cargo genes, the PIs may exhibit characteristics of clustered prophage genes accompanied by scattered bacterial elements (Fig. 1a). Here, we cluster adjacent prophage ORFs into a single PI, ensuring that the intergenic distance between adjacent prophage ORFs is less than *D*. And to obtain relatively high-confidence PIs, we ensure that the PI score (*P*), which is the average probability of each ORF based on the MLP output, is not less than 0.7. If the *P* falls below 0.7, we iteratively trim ORFs one by one from both ends until *P* reaches or exceeds 0.7. Empirically, we require the PI to include at least five phage ORFs to output PI information(21) (Fig. 1b).

**Fig. 1.**
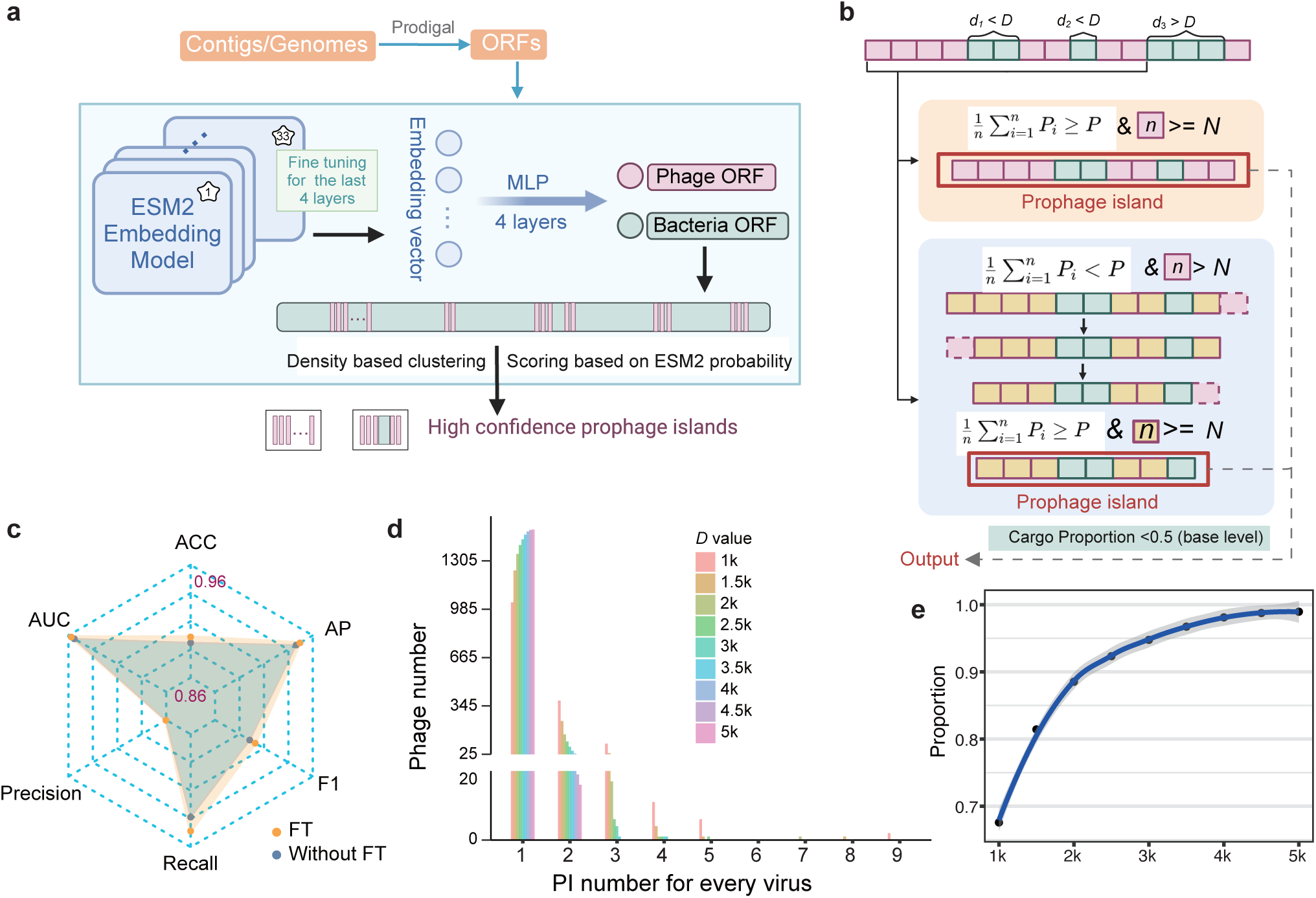
The PIDE framework for identifying PIs. **a**, Schematic overview of PIDE. PIDE takes DNA sequences as input and outputs the predicted PI regions. **b**, The detailed strategy employed by PIDE for clustering prophage ORFs and identifying prophage regions. *D* represents the distance, and it can vary depending on different requirements. **c**, The performance of PIDE and the effect of ablation analysis. FT: fine-tuning. **d**, The distribution of PIDE-predicted PIs across phage genomes under different *D*. The x-axis represents the number of PIs predicted by PIDE for each complete phage genome, and the y-axis indicates the number of complete phage genomes. **e**, The x-axis represents different *D* values, and the y-axis indicates the proportion of complete phages that can be recovered by PIDE.

The accuracy of the model on the test set reached 0.90, the F1 score reached 0.90, the AUC reached 0.96, and the AP reached 0.95 (Fig. 1c, Table 1). To investigate the contribution of the step of finetuning ESM-2, we performed an ablation study. Without finetuning, the evaluation performance of the model dropped, especially in the recall, demonstrating the benefit of fine-tuning (Fig. 1c). Overall, the model demonstrates strong proficiency in identifying phage genes.

**Table 1.** The model performance.

|  | ACC | AUC | Precision | Recall | F1 | AP |
| --- | --- | --- | --- | --- | --- | --- |
| Without_FT | 0.89 | 0.95 | 0.86 | 0.93 | 0.90 | 0.94 |
| FT | 0.90 | 0.96 | 0.86 | 0.95 | 0.90 | 0.95 |

### Determination of the clustering distance for PI identification

Next, we assessed the impact of different *D* values on PI identification, as larger *D* values tend to introduce false positives by including bacterial sequences within prophages, whereas smaller *D* values can fragment prophage sequences, leading to the omission of potential prophage-associated genes. To assess the effectiveness of different *D* values in predicting prophages, we calculated the number of PIs predicted by PIDE for each single complete PhaTYP-predicted temperate phage genome from the RefSeq(22,23). A greater number of predicted PIs within a single complete temperate phage suggests lower effectiveness of PI prediction. Nine different *D* values ranging from 1 kb to 5 kb were tested. At the distance threshold of 1 kb, PIDE fragments 32% of phages into two or even nine segments; at 3 kb, 95% of phages remain unfragmented. And at 5 kb, 99% remain unfragmented (Fig. 1d, e). Here, PIDE defaults to the 3 kb threshold. At this threshold, further analysis revealed that 93.5% of PIs predicted by PIDE achieve a genome coverage of over 90% for the original reference genome, with the missing regions mainly located at the ends of the phage genome (Additional file 1: Fig. S1a, S1b).

### PIDE enables the discovery of extensive PIs with precise boundary detection

We evaluated PIDE alongside three other widely-used tools on 38 gut-derived bacterial isolates with complete whole genome sequences available. A total of 365 PIs were identified using PIDE, whereas geNomad, Virsorter2, and PHASTER identified 142, 101, and 174 PIs, respectively. PIDE successfully predicted 84% of geNomad predictions, 80% of Virsorter2 predictions, and 55% of PHASTER predictions, demonstrating the highest performance in recovering overlapping PIs across different tools (Fig. 2a). Moreover, PIDE identified the highest number of unique PIs not predicted by other tools, with 184 out of 365 PIs being unique, followed by PHASTER (Additional file 1: Fig. S2a). To validate these predicted PIs, we conducted BLAST searches against the NT database (evalue < 1E-10) to identify potential viral sequences which have been previously reported. Among the 181 PIs predicted by PIDE and at least one other tool, 132 were alignable, with 99 exhibiting coverage of ≥ 50% (Fig. 2b). For unique PIs with coverage ≥ 50%, PIDE had the highest alignable count at 24, followed by PHASTER with 5, VirSorter with 1, and geNomad with 0 (Fig. 2c, Additional file 1: Fig. S2a). The remaining unalignable sequences may represent novel, unidentified phages, although they could also include false positives. Overall, these results demonstrated the superior sensitivity of PIDE in PI prediction.

**Fig. 2.**
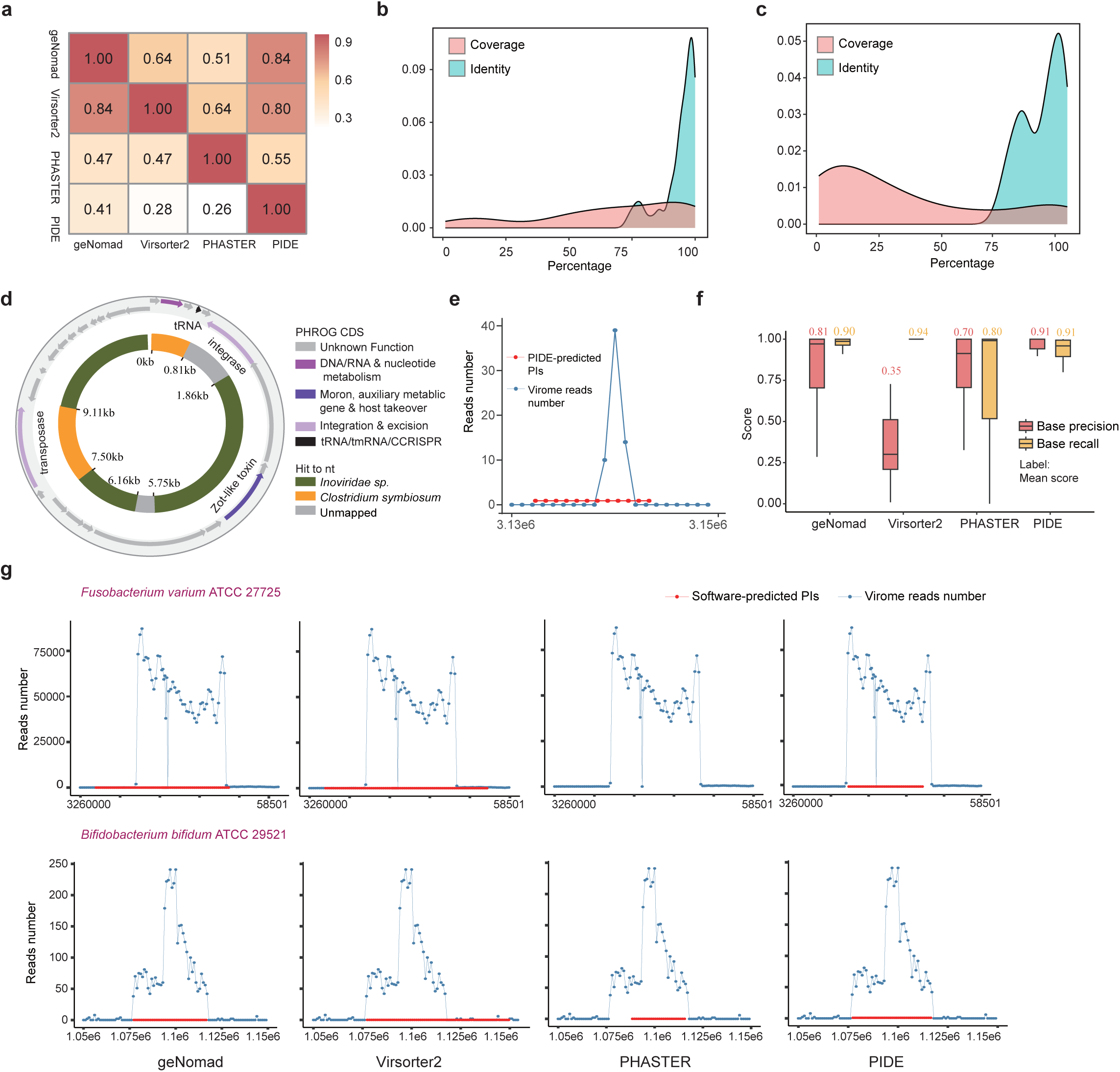
The performance of PIDE and other tools in PI detection. **a,** The heatmap showing the extent of shared PIs predicted by different tools. Each row represents a tool, with cells showing the proportion of PIs predicted by other tools relative to the total PIs predicted by that tool. Density plots showing the distribution of coverage and identity for 132 overlapped-predicted PIs **(b)** and 89 unique-predicted PIs **(c)** aligned with virus entries of nt database. **d,** Gene annotations of PIDE-Csy PI (outer ring) and the alignments with the nt database (inner ring). **e,** The distribution of virome reads on local *Clostridium symbiosum* ATCC14940 (blue), with PIDE-Csy PI region highlighted in red. **f,** The boxplot shows the distribution of precision and recall for PIs predicted by different tools, measured at the base level. **g,** The distribution of virome reads on local *Fusobacterium varium* ATCC 27725 and *Bifidobacterium bifidum* ATCC 29521 (blue), with PI regions predicted by different tools highlighted in red.

We then selected a PI, named PIDE-Csy PI, exclusively detected by PIDE, originating from *Clostridium symbiosum* ATCC14940 and measuring 11,607 bp in length, for further analysis. Genome annotation using Pharokka identified an integrase gene, which is a characteristic element of phages, as well as a Zonula Occludens Toxin (Zot)-like toxin gene(24). Subsequent BLASTn analysis in the NCBI database revealed that this sequence shares over 97% similarity and 66% coverage with *Inoviridae sp*. isolate ctnAm19 (Fig. 2d). Additionally, sequencing data of viral-like particles (VLPs) from this strain, treated with a prophage induction inducer, indicated discernible activity of the PIDE-Csy PI. Given that Zot is an enterotoxin capable of reversibly altering the tight junctions of intestinal epithelia(25), the status of this PI could potentially affect gut health (Fig. 2e).

The VLP data can help provide precise insights into the boundaries of active PIs, which can serve as a reliable ground truth. Hence, we computed and compared the base recall and base precision metrics for PIs identified by different tools (See Methods), aiming to assess their accuracy in boundary delineation. The results demonstrated that PIDE exhibited the highest base precision with a mean score of 0.91, followed by geNomad at 0.81, PHASTER at 0.7, and Virsorter2 at 0.35. In terms of base recall, Virsorter2 achieved the highest mean score at 0.94, followed by PIDE at 0.91, geNomad at 0.90, and PHASTER at 0.80 (Fig. 2f, g). Overall, these results indicate that PIDE outperforms other tools in independent validation tests, enhancing our capacity to infer the gene pool and ecological dynamics of naturally occurring PIs.

### PIDE facilitates detection of phage contigs across diverse sequence lengths

PIDE can also be used to identify phage-derived contigs. To determine whether a contig is phage-derived, we assessed the proportion of phage-like ORFs relative to the total number of ORFs. Contig length can affect the accuracy of identifying phage contigs, so we first established the optimal thresholds for the phage-like ORF proportion across different contig lengths. We simulated 60,000 contigs of various lengths (0.5 kb, 1 kb, 3 kb, 5 kb, 10 kb, 20 kb) from 6,099 phage genomes in RefSeq and 4,744 representative bacterial genomes from the Unified Human Gastrointestinal Genome (UHGG). These simulated contigs were analyzed by PIDE for phage ORF prediction, and the phage-like ORF proportions were calculated. The optimal thresholds for contigs of 0.5 kb, 1 kb, 3 kb, 5 kb, 10 kb, and 20 kb were determined to be 1, 1, 0.57, 0.56, 0.53, and 0.52, respectively (Fig. 3a).

**Fig. 3.**
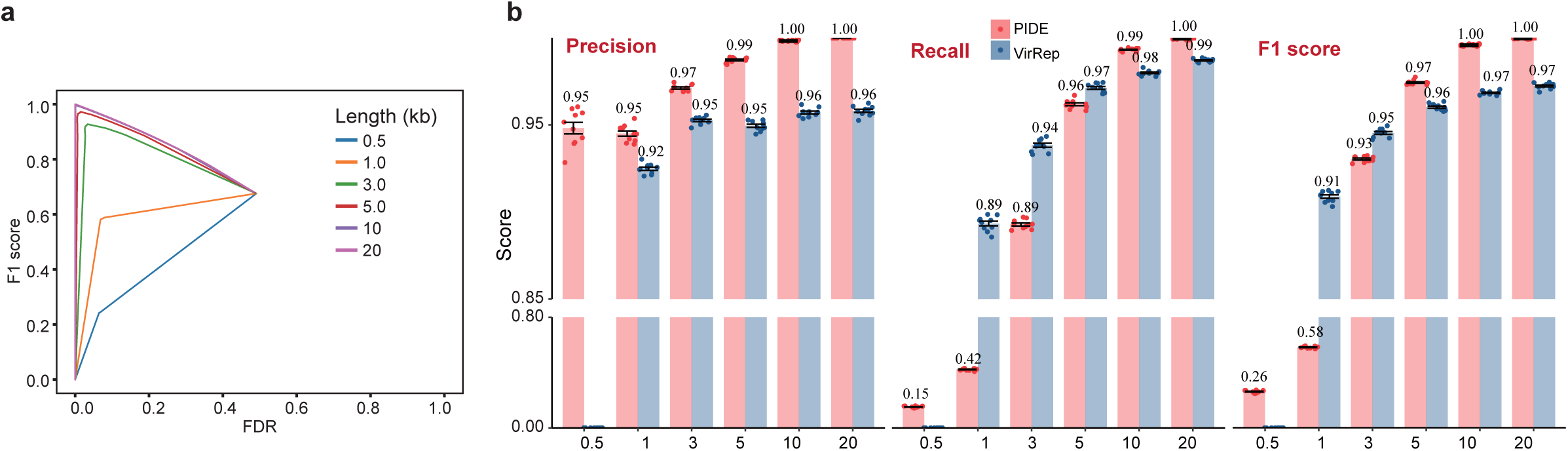
The performance of PIDE in identifying assembled phage contigs. **a,** F1 score versus FDR curve at various sequence lengths. **b,** The sequence classification performance of PIDE and VirRep across sequences of varying length. Performance was measured using precision, recall, and F1 score.

Performance metrics were assessed using an additional set of ten independent samples and compared with VirRep, a state-of-the-art hybrid language representation learning framework for identifying viruses from metagenomic contigs(27). PIDE consistently exhibited higher precision than VirRep across all sequence lengths (Fig. 3b). Unlike VirRep, which is limited to sequences longer than 1 kb, PIDE could also be applicable to shorter sequences (0.5-1 kb). However, in this shorter length range, PIDE detected less than 50% of phage sequences. F1 score comparisons indicated that PIDE showed an advantage over VirRep in predicting phage contigs for longer sequences, except for sequences under 3k (Fig. 3b).

### The features of PIs in the human gut microbiome

Next, we applied PIDE to 4,744 species representatives derived from the human gut microbiome to survey the PI features in the human gut microbiome. A total of 24,467 PIs were detected across 4,198 genomes (88.5%). These islands varied widely in size, with a median of 7738 bp, and up to 205,026 bp (Additional file 1: Fig. S3a). On average, prophage-derived sequences constituted 2.8% of their host genomic content, with an interquartile range from 0.8% to 3.6%. A genome contained an average of 6 PIs, with a maximum of 42. Among the four most prevalent phyla and genera, Actinobacteriota genomes exhibited lower PI counts compared to Proteobacteria genomes, which typically harbored higher numbers of PIs. At the genus level, *Collinsella* showed the lowest relative abundance of PIs (Fig. 4a,b). This differential distribution suggests the occurrence frequency of lysogeny varies significantly among different taxa.

**Fig. 4.**
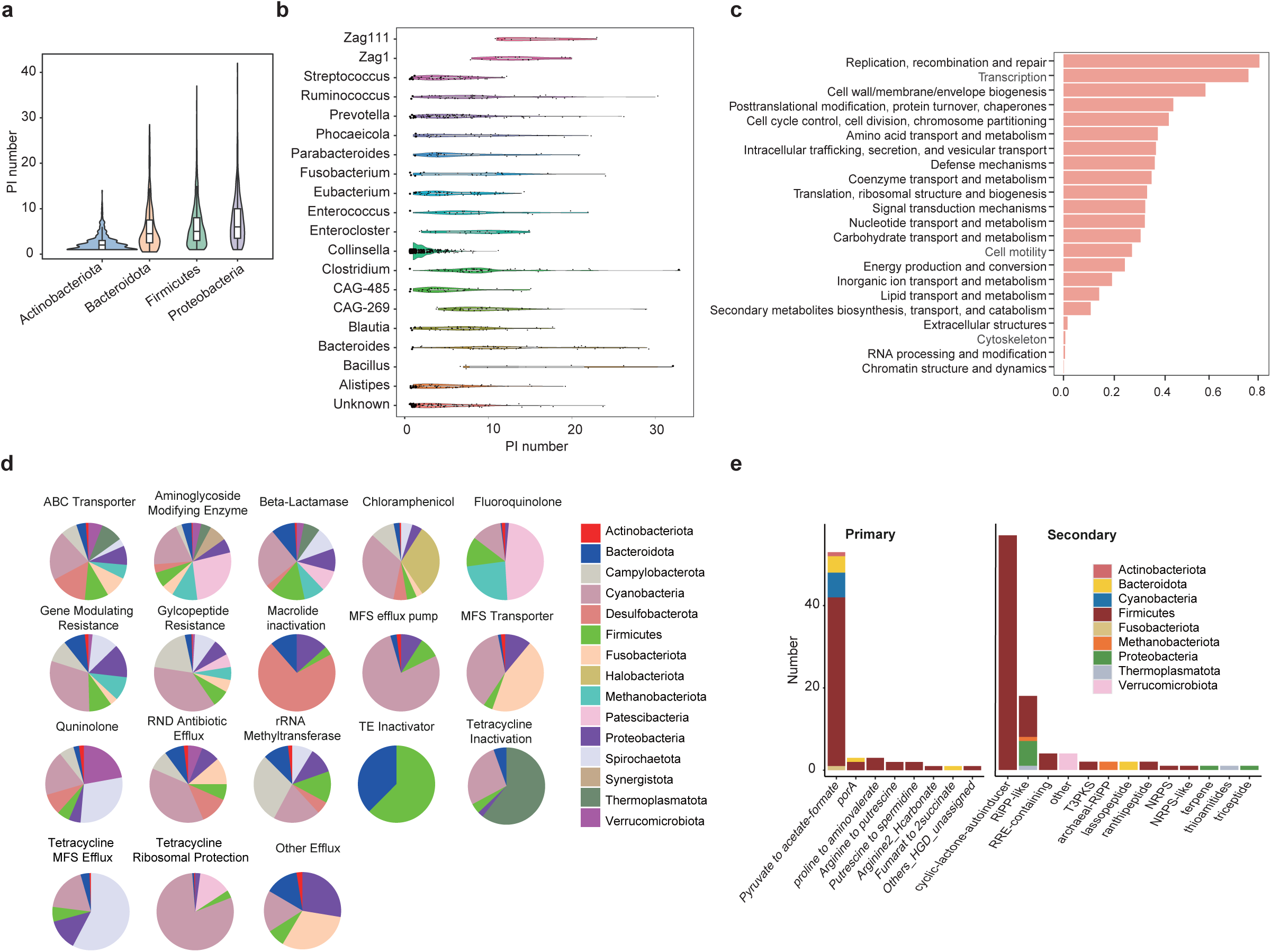
The application of PIDE to 4,744 representative prokaryotic genomes from the human gut microbiome. The violin plot displays the distribution of the number of PIs on each genome across the top four most abundant phyla **(a)** or the top twenty most abundant genera **(b)**. **c,** eggNOG annotation of PIs, where the x-axis represents the prevalence in prokaryotic genomes and y-axis represents different eggNOG categories. **d,** The relative abundance of PI-encoded ARGs across different phyla within various categories. **e, The** distribution of the number of PI-encoded genes across different phyla in various metabolic pathways.

### PIs carry a wide repertoire of cargo genes

Among these 4,198 lysogens, 4,039 (96.2%) were predicted to containing PIs carrying cargo genes, with the proportion of cargo genes within the PIs ranging from 0% to 87.7% (Additional file 1: Fig. S3b). To understand the potential functions of these cargo genes, we first annotated PI ORFs using eggNOG-mapper(28). And 183,410 out of 399,565 (45.9%) were assigned to the specific COG categories. Then we calculated the prevalence of each COG category encoded by PIs in all bacterial genomes. In addition to genes typically found in phages related to replication, recombination, and transcription, a significant proportion of bacterial PIs also carry genes involved in regulating the bacterial cell cycle, amino acid transport, and metabolism, as well as defense-related pathways, including genes containing ParBc, AAA_31, ParE_toxin, ABC_tran domains (Fig. 4c).

Due to the potential horizontal transfer of antibiotic resistance genes (ARGs) from commensal bacteria in the gut, especially those within PIs, we conducted a more detailed analysis of the ARGs carried on PIs. Our analysis showed that 34.2% of bacteria contained PIs carrying 120 different types of ARGs categorized into 18 types involving resistance to various antibiotics through different mechanisms (Additional file 1: Fig. S3c). The most abundant types of ARGs included genes resistant to a wide range of antibiotics, such as beta-lactamases, glycopeptides, aminoglycosides, chloramphenicol, fluoroquinolones, and tetracyclines. Next, we normalized for the absolute number of bacteria within each phylum to compare the proportions of PIs carrying different types of ARGs across various phyla. The result indicated PIs of Cyanobacteria nearly carried all kinds of ARGs, except for TE inactivator (Fig. 4d). Previous studies have revealed Cyanobacteria serve as vital reservoirs for antibiotic resistance genes(29). Apart from these bacteria that may originate from environmental contamination, PIs carrying ARGs were mainly from Bacteroidetes, Proteobacteria, and Firmicutes, while they occured less frequently in Actinobacteria.

Moreover, we found that 3.3% of bacteria carried PIs harboring metabolic related genes, which might modulate bacteria physiology. Pyruvate to acetate/formate and cyclic-lactone-autoinducer were the most abundant primary and secondary pathway, respectively (Fig. 4e). And the main PI carriers of these related genes were from Firmicutes. Overall, these results demonstrate that the PIs harbor a diverse array of functional genes, which may contribute to the physiological processes within the human gut microbiota.

## Discussion

Identifying prophages is essential for elucidating the mechanisms of bacteria-phage interactions and understanding microbial community structure and function(30). Here, we introduce PIDE, a novel framework for identifying prophages in whole bacterial genomes or MAGs. Large pretrained protein language models have demonstrated significant progress in annotating protein functions(18–20). As a tool utilizing a large language model, PIDE performs exceptionally well in distinguishing phage genes from bacterial genes, a crucial step in recovering PIs that may have been previously overlooked. By sequencing phages induced from isolated bacterial strains, we have generated reliable data that showcases active prophage regions, providing a benchmark for evaluating the performance of various tools. Compared to previous tools, the boundaries predicted by PIDE can achieve both high base recall and base precision at the nucleotide level. To be noted, PIDE identifies better prophage boundaries by accurately identifying the first or last phage ORF of the prophage, which can narrow down the search range for these attachment sites. However, it cannot predict the exact attachment regions of prophages. Other methods, identifying the split reads generated by lysis-lysogeny switch or searching for attachment site core sequences, such as Prophage tracer (13,31), can be integrated for further identification of the prophage attachment sites.

PIDE is also capable of predicting phage contigs in metagenomic data, even though it was not originally designed for this purpose. Considering that it is based on identifying multiple prophage genes within a certain region, it is understandable that precise results cannot be obtained when predicting shorter contigs (<= 3kb). Another algorithm, which can be used to predict short viral contig accurately, such as VirRep(27), can be integrated with PIDEI for a good analysis of viral contig searching.

Applying PIDE to human gut microbiome revealed the widespread presence of prophages, as 88.5% of the bacteria harbor PIs. Furthermore, these PIs harbor various genes associated with bacterial survival and adaptation, including antibiotic-resistance genes and metabolism-related genes. The identified prophage-encoded genes could enhance our understanding of bacterial ecology and evolution, aid in the development of novel antimicrobial strategies, and inform the design of targeted therapeutic interventions(30,32–34); further experiments are needed to elucidate the functions of prophage-encoded genes.

## Conclusions

In conclusion, PIDE offers a highly accurate method for identifying prophage island regions with precise boundary detection, utilizing both bacterial and metagenome-assembled genomes. Additionally, it can be employed for viral contig identification. We believe PIDE will enhance the exploration of microbiome and virome data, contributing valuable insights to microbiome research.

## Methods

### Dataset generation

The phage and prokaryotic proteins from UniRef50 database (May 2024 version) were downloaded for training the classification model. The phage dataset comprised 263,843 sequences, while the bacteria and archaea dataset contained 40,572,403 sequences. Highly similar sequences to those in the phage dataset were removed from the bacteria and archaea datasets with cd-hit-2d (sequence similarity>95%)(35), resulting in 40,566,921 sequences. Subsequently, 263,843 sequences were randomly sampled from these 40,566,921 sequences to achieve a 1:1 ratio of positive to negative samples. Then the dataset was randomly partitioned into training, validation, and test datasets with a ratio of 7:1.5:1.5.

### Model training processes and test metrics

The ESM-2 model with 33 Transformer Encoder layers and 650M parameters were used for the classification of phage protein. The input protein sequence is first tokenized and subsequently encoded into an embedding representation. Then the embeddings are averaged based on sequence length, yielding a 1280-dimensional vector. This vector is fed into a five-layer MLP, where each layer employs ReLU activation. Finally, the probability value of phage protein is obtained by processing the MLP output using a softmax layer. A protein with a value greater than 0.5 is generally considered to be a phage protein. During training, the last four layers of the Transformer Encoder were fine-tuned, while the remaining layers were kept frozen. Additionally, the MLP parameters were optimized from scratch. The learning rate was set to 5e-6, with a batch size of 4 and a weight decay of 1e-5. The model converged after 2 epochs. A variety of metrics were chosen to evaluate and compare the performance of models. For the binary task, we calculated the Accuracy (ACC), Precision, Recall, F1, Area Under Curve (AUC), and Average Precision (AP) value. The formulas for ACC, Precision, Recall, and F1 are as follows (1)-(4):

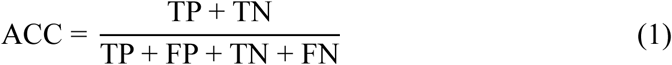

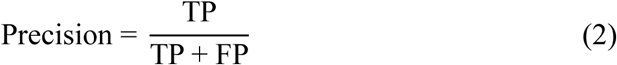

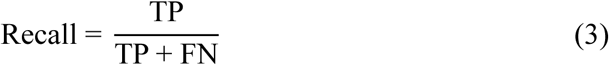

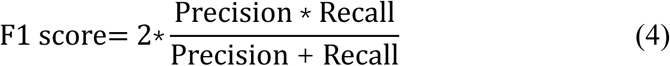

where TP, FP, TN, and FN mean the number of true positive, false positive, true negative, and false negative samples during a test.

### Performance benchmarks

Prophages can enter lytic induction spontaneously or upon induction by mitomycin C or other agents, initiating replication, proliferation, and the release of numerous progenies from bacteria(30). As a result, the majority of reads from VLPs originate from potentially active prophages, leading to significantly higher read coverage in these regions compared to the surrounding regions. Given that there are few data sets that accurately annotate the prophage region of bacteria and lack a gold standard for evaluating the effectiveness of different tools, we isolated and purified VLPs from 38 different bacterial culture. The VLP DNA was extracted and amplified for sequencing.

The raw sequencing reads were first treated using fastp to remove adapters, duplicate reads and low-complexity reads(36). Then the clean reads were mapped against the bacterial genome using bowtie2(37). To pinpoint these active regions, Homer (-style histone) was applied to call peaks, particularly adept at identifying broad peaks(38). Given that most prophages span several to tens of kilobases and there is a feature of non-uniform read coverage across prophage positions, peaks within the same prophage may appear discontinuous (Additional file 1: Fig. S2b). Bedtools was employed to merge peaks within 6 kb into a single peak to reconcile this(39).

The following tools were included in our benchmarks: geNomad (version 1.8.0), VirSorter2 (version 2.2.4), and PHASTER. The tools were executed with default parameters.

### Comparison of base recall and base precision among different tools

Given that PIs are cryptic and some others could not be triggered, we focused our comparison on the potential inducible prophage regions which were defined as the union of prophages predicted by the four tools that intersected with the identified peaks. And the regions of these peaks were used as the experimental gold standard to evaluate the performance of the four tools in predicting prophages. For each peak, the base recall score was defined as the proportion of the region predicted by a specific tool that covered the peak region. For each predicted prophage by a specific tool, the base precision score was defined as the proportion of the peak region that covered the predicted prophage region.

### Applying PIDE to the human gut microbiome

PIED was employed to analyze the prophage characteristics in the human gut microbiome. 4,744 representative genomes of species derived from the human gut microbiome were retrieved from http://ftp.ebi.ac.uk/pub/databases/metagenomics/mgnify_genomes/human-gut/v2.0.2/species_catalogue/ (26).

The PI sequences were extracted using seqtk (https://github.com/lh3/seqtk) and functionally annotated using eggNOG-mapper with default settings(28). gutSMASH and antiSMASH were employed to annotate genes involved in primary and secondary metabolism on prophages, respectively(40,41). And these PIs were also queried against the Resfams profile HMM database to identify potential resistance genes(42).

## Declarations

### Ethics approval and consent to participate

Not applicable.

### Consent for publication

Not applicable.

### Availability of data and materials

All datasets used in this study can be retrieved from the following links: (1) Uniprot, https://www.uniprot.org; (2) Refseq,

https://ftp.ncbi.nlm.nih.gov/refseq/release/viral/viral.1.1.genomic.fna.gz; (3) UHGG, http://ftp.ebi.ac.uk/pub/databases/metagenomics/mgnify_genomes/; The parameter of the model can be downloaded from

https://zenodo.org/records/12759619/files/PIDE.model.tar.gz?download=1. The source code for PIDE has been placed into the GitHub

(https://github.com/chyghy/PIDE ). All the data are available from the corresponding author on reasonable request.

### Authors’ contributions

H.G., B.L. and G.L. conceived the project and wrote the manuscript. B.L. and H.G. designed the method. Z.G. conducted experiments. Z.G. and L.Z. helped with data analysis. All authors revised the manuscript and approved the final version for submission.

## Acknowledgments

We are grateful to members of the Liang laboratory for their help and suggestions.

## Additional files

Additional file 1.pdf: Supplementary figures, Fig. S1-S3

