## Supplementary material for "Highly accurate prophage island detection with PIDE": Fig. S1, Fig. S2, Fig. S3

### Supplementary figures

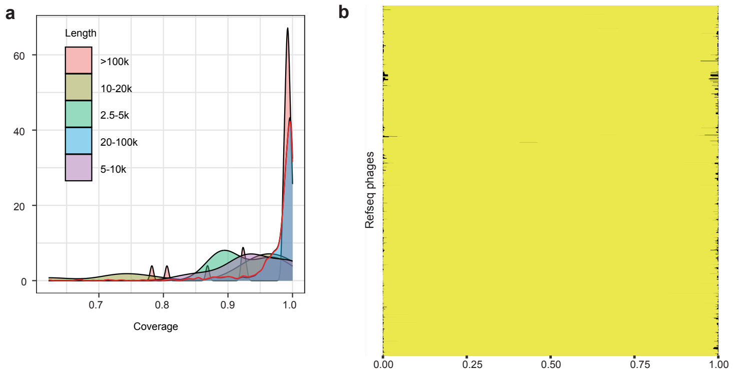


**Fig. S1 The ability of PIDE to recover phage genomes with *D* = 3kb. a,** Coverage distribution of PIDE-predicted PIs in original phage genomes across various length ranges. Red line represents the overall distribution. **b,** The coverage of PIDE-predicted PIs on the original phage sequences. All sequence lengths are normalized to 1. Each horizontal line represents an original phage sequence, with yellow color indicating the regions covered by PIs and black indicating the PI-uncovered regions.

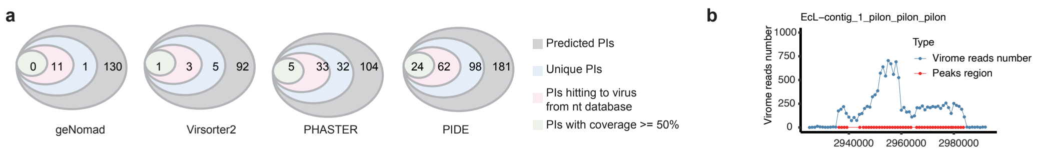


**Fig. S2 PIs predicted by different tools. a,** Plot indicating the total number of PIs predicted by each of the four software tools, the number of unique PIs for each software, the number of unique PIs matched to viruses in the nr database, and the number of matches with coverage ≥50%. **b,** An example showing the regions of Homer-predicted peaks and the distribution of virome reads in those regions and their surrounding areas.

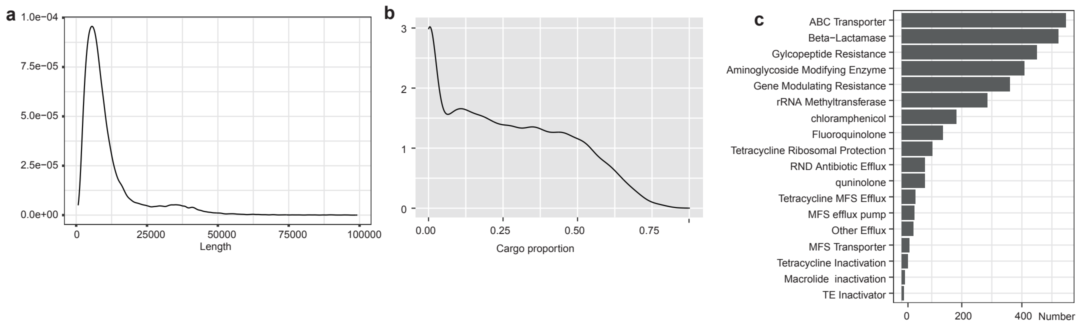


**Fig. S3 The characteristics of PIs in human gut microbiome. a,** The length distribution of the 24,467 PIs. **b,** The distribution of the proportion of cargo genes relative to the total genes in PIs. **c,** The number of PIs carrying different types of ARGs.
